# High-resolution spatial transcriptomics maps viral tropism and reveals spatially organized immune modules in viral encephalitis

**DOI:** 10.64898/2026.08.17.745247

**Authors:** Chase Holdener, Shaowen Jiang, Kira A. Griswold, Kayla M. Kimble, Peter Schweitzer, Madhav Mantri, Meleana Hinchman, Danica M. Sutherland, John Parker, Terence S. Dermody, Iwijn De Vlaminck

**Affiliations:** Meinig School of Biomedical Engineering, Cornell University, Ithaca, New York, USA; Department of Computational Biology, Cornell University, Ithaca, New York, USA; Department of Microbiology and Molecular Genetics, University of Pittsburgh School of Medicine, Pittsburgh, Pennsylvania, USA; Institute of Infection, Inflammation, and Immunity, UPMC Children’s Hospital of Pittsburgh, Pittsburgh, Pennsylvania, USA; Baker Institute for Animal Health, College of Veterinary Medicine, Cornell University, Ithaca, New York, USA; Department of Pediatrics, University of Pittsburgh School of Medicine, Pittsburgh, Pennsylvania, USA

**Keywords:** Viral encephalitis, viral tropism, reovirus, spatial transcriptomics, co-expression networks, neural antiviral immune responses

## Abstract

Viral encephalitis is a debilitating disease that most commonly affects vulnerable populations, including the very young and elderly. Despite its severity, few countermeasures exist due to the structural and immunological complexities of the central nervous system (CNS). To better understand the spatial dynamics of viral spread and the concurrent host immune responses, we used high-resolution spatial transcriptomics to profile reovirus infection in the neonatal mouse brain at 3, 5, and 7 days post-infection. We constructed a comprehensive spatiotemporal atlas of infection, which revealed viral dissemination from sites of cerebrospinal fluid circulation to neighboring brain parenchyma, with enriched infection in the thalamus and midbrain coincident with *Slc17a6* (VGLUT2) excitatory neurons. Unbiased spatial gene co-expression network analysis uncovered rich gene lists linked to temporal waves of host immune responses, originating with interferon-stimulated gene modules, followed by myeloid cell infiltration and adaptive cytotoxic T-cell responses at times of peak disease. Furthermore, we identified spatial correlation between viral transcripts and several upregulated host snoRNA-related genes (e.g., *Nop58* and *Snhg1*) with high-confidence, suggesting viral use of host ribosomal modification machinery. We also observed a strong anti-correlation of astrocyte markers with reovirus transcripts, suggesting glial cell disruption. This work provides a high-definition spatial framework to understand interactions between viral infection and host immunity in the brain.

## Introduction

Viral pathogens of the central nervous system (CNS) are a significant cause of morbidity and mortality^1–3^. This burden is partly due to diagnostic and therapeutic gaps, as viral origins remain unidentified in up to 60% of cases of presumed viral encephalitis^4^. Viral encephalitis occurs following either hematogenous spread of a neurotropic virus to the CNS or neural spread within peripheral axons^5,6^. New insights into viral and host factors responsible for viral neurotropism, dissemination within the CNS, and innate and adaptive immune responses are required to fully understand mechanisms of viral CNS disease, develop effective countermeasures, and enable engineering of highly selective vectors that infect discrete regions in the brain for use as oncolytics and other targeted therapeutics.

Mammalian orthoreoviruses (colloquially called reoviruses) are nonenveloped, double-stranded RNA viruses that can infect the CNS and cause lethal meningoencephalitis in newborn mice^7^. The age-dependent susceptibility to reovirus neurologic disease^8^ makes reovirus a useful experimental system for studying mechanisms of viral neuropathogenesis and the antiviral immune response in the developing brain. While the effects of reovirus infection are documented^9–11^, mechanisms by which the virus navigates the complex architecture of the brain, traversing specific barriers and targeting distinct neuronal subpopulations, are not well understood.

Our understanding of reovirus encephalitis largely has relied on viral load measurements and traditional histology^12,13^, which assess viral replication efficiency and tissue morphology but lack unbiased, high-dimensional gene expression sampling in areas of the brain infected by the virus. Bulk RNA sequencing provides insights into transcriptional programs but sacrifices the spatial context required to define the infected microenvironment from bystander tissues^14^. Thus, the complex heterogeneity of the brain, comprising diverse neuronal subtypes, supporting cells, and vascular components, requires a methodology that captures spatial context while providing rich gene expression data.

In this study, we used the Slide-seq V2 high-resolution spatial transcriptomics platform^15^ to produce a spatiotemporal map of reovirus infection in the neonatal mouse brain over a time course. Achieving near single-cell spatial resolution, we obtained high-quality gene expression data from brain regions and cell types that enabled us to follow reovirus spread from the ventricular system into the parenchyma. Using virus and host gene spatial co-expression network analyses, we identified over 100 spatial gene modules in the brain, including over 10 dynamic immune-related or virus-associated modules that are activated or disrupted only in infected brains. Temporal gene expression analyses uncovered waves of distinct, spatially defined facets of the host immune response. Finally, we observed new virus-host interactions, including a link between viral load and upregulated host snoRNA-related genes, providing new insights into the molecular architecture of viral encephalitis.

## Results

### High-resolution spatial transcriptomic profiling of reovirus-infected mouse brains

To study the pathogenesis of reovirus encephalitis, we first used high-resolution spatial transcriptomics to define transcriptional changes in brain tissue during disease progression. Neonatal mice were inoculated intracranially in the right-brain hemisphere with reovirus or vehicle control (mock), euthanized at 3, 5, or 7 days post-inoculation (DPI), representing early, middle, and late stages of disease^16–18^, and brain tissue was collected. Left brain hemispheres were used to quantify viral titers, and right brain hemispheres were frozen and processed for immunohistochemistry, fluorescence in-situ hybridization (FISH), and spatial transcriptomics using the Curio Seeker platform (**Fig. 1a**). Spatial transcriptomics sequence reads were aligned to the mouse mm39 and reovirus genomes, and after data quality control and filtering, we obtained 1,155,702 high-resolution spatial transcriptomes across the five samples (**Supplementary Table S1**). There are two biological replicates for 7 DPI referred to as 7 DPI (rep. 1) and 7 DPI (rep. 2). This extensive dataset provides near single-cell resolution (10 μm) spatial profiling of the infected brain microenvironment throughout reovirus infection progression.

**Fig. 1.**
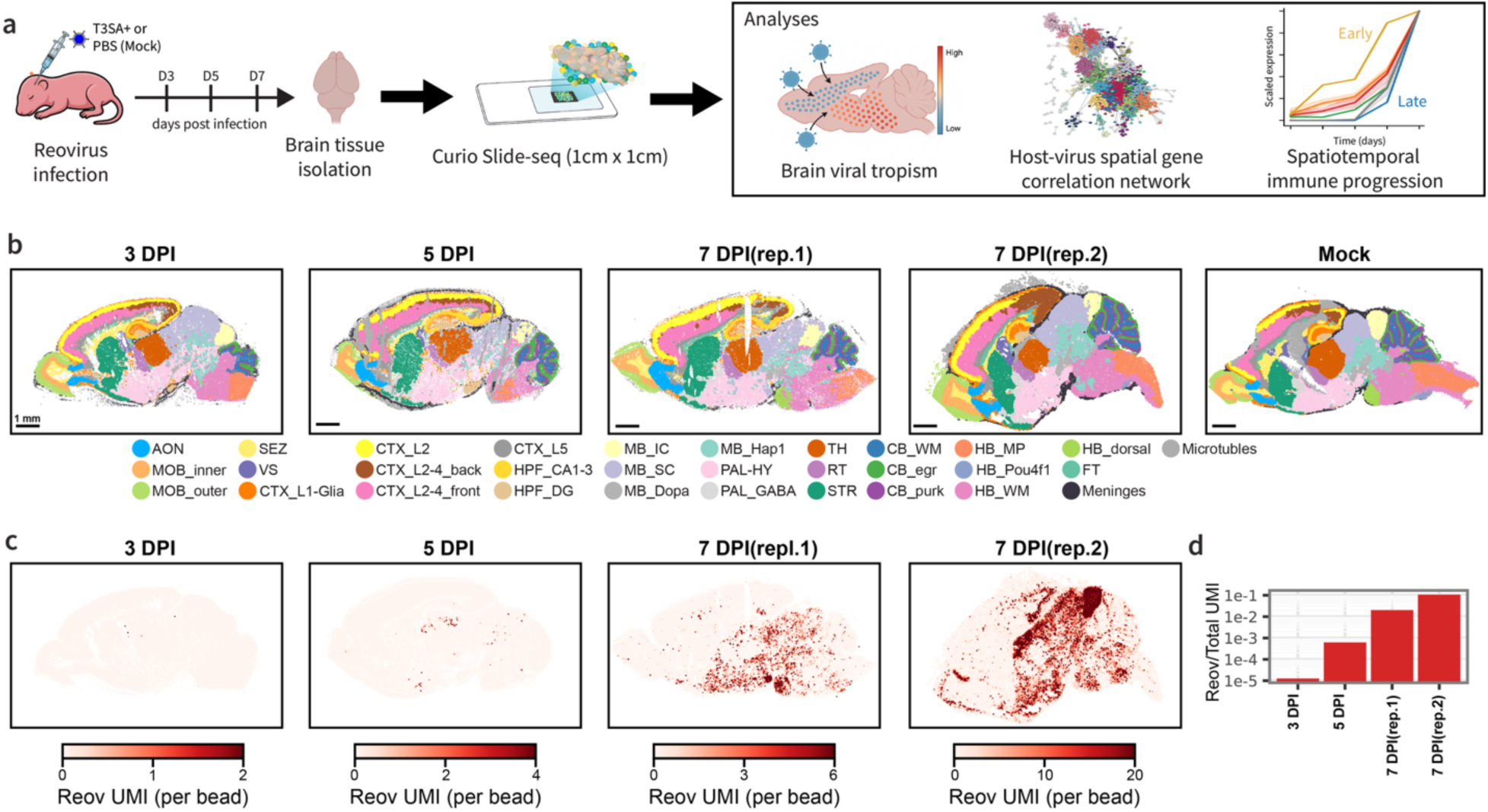
Spatial transcriptomics of brain tissues of reovirus-infected neonatal mice. **a** Experimental design and analysis workflow. Three-day-old mice were inoculated intracranially with reovirus T3SA+ or PBS control in the left hemisphere. Brain tissues were collected at 3, 5, or 7 days post-infection. Right brain hemispheres were processed for spatial transcriptomics. Left brain hemisphere were process for viral plaque assay. Computational analyses included brain region and cell type annotation, viral tropism determination, virus-host spatial gene correlation network analysis, and spatiotemporal immune response characterization. **b** Brain region annotations across infection time course samples and mock control. Scale bars, 1 mm. **c** Spatial mapping of reovirus transcripts (Reov UMI) during the infection time course. **d** Total reovirus UMI counts during infection time course.

The poly(A)-enriched spatial transcriptomics captured substantial amounts of reovirus RNA (**Fig. 1c**) despite the reovirus genome and mRNAs lacking poly(A) tails^19^. Total reovirus RNA counts increased progressively from 3 to 7 DPI (**Fig. 1d**) and matched trends observed for viral titers quantified from contralateral hemispheres (**Fig. 1e**). We captured transcripts from all 10 reovirus gene segments and did not observe any relationship between virus transcript abundance and gene segment length or poly-U stretch frequency (**Supplementary Fig. 2a**). For subsequent analyses, we annotated regional and cell type RNAs for each of the 5 spatial datasets. Using GraphST^20^, we resolved 32 distinct anatomical subregions (**Fig. 1b, Supp. Fig. 1c**) organized into 12 major brain structures (**Supp. Fig. 1a,b**), including olfactory areas (OLF), ventricular systems (VS), cerebral cortex layers (CTX), hippocampal formation (HPF), striatum (STR), pallidum-hypothalamus (PAL-HY), thalamus (TH), midbrain (MB), cerebellum (CB), hindbrain (HB), fiber tracts (FT), and meninges. (**Fig. 1b and Supplementary Fig. 1a,b**). For cell type annotation, we conducted standard dimensionality reduction and clustering, followed by manual annotation with cross-referencing marker genes on established cell signatures from the Allen Mouse Brain Atlas^21^. This analysis yielded 35 distinct cell-type labels, including different neuronal subtypes, glial cells (astrocytes, oligodendrocytes, and microglia), vascular cells, choroid plexus epithelial cells, and others (**Supplementary Fig. 1d,e**). To extend these annotations across all samples, we used a spatial-to-spatial deconvolution strategy using PrismST^22^. Rather than relying on external single-cell RNA-seq references, which can introduce platform-specific biases, we used our deeply sequenced spatial datasets, Mock and 7 DPI (rep. 2), as internal references. This approach preserved cell type-specific expression signatures and enabled robust cell type assignment across samples with varying sequencing depths. Overall, our regional and cellular annotations are consistent with the known, complex architecture of the brain^21^.

### Spatiotemporal distribution of reovirus transcripts reveals potential pathways of viral dissemination and tropism

Reovirus disseminates by both neural and hematogenous routes to reach the brain^13,23^ and targets specific subsets of neurons for infection^24^. However, the exact mechanisms of these processes are poorly understood. To investigate reovirus tropism, we plotted the spatial distribution of reovirus transcripts across timepoints in our collected datasets. At 3 DPI, reovirus transcripts were sparse but appeared associated with the ventricular system (**Fig. 2a**), which produces and circulates cerebrospinal fluid (CSF) throughout the CNS. Indirect immunofluorescence experiments confirmed sites of reovirus replication in ventricular tissue (**Fig. 2b**). By 5 and 7 DPI, viral transcript distribution shifted from the ventricular system to the brain parenchyma (**Fig. 2a**). The thalamus and midbrain were major sites of infection at these time points, with viral transcripts accumulating over time in these regions. Collectively, this ventricle-to-parenchyma spread suggests that reovirus uses CSF circulation through the ventricles and meninges to gain access to susceptible neuronal populations throughout the brain.

**Fig. 2.**
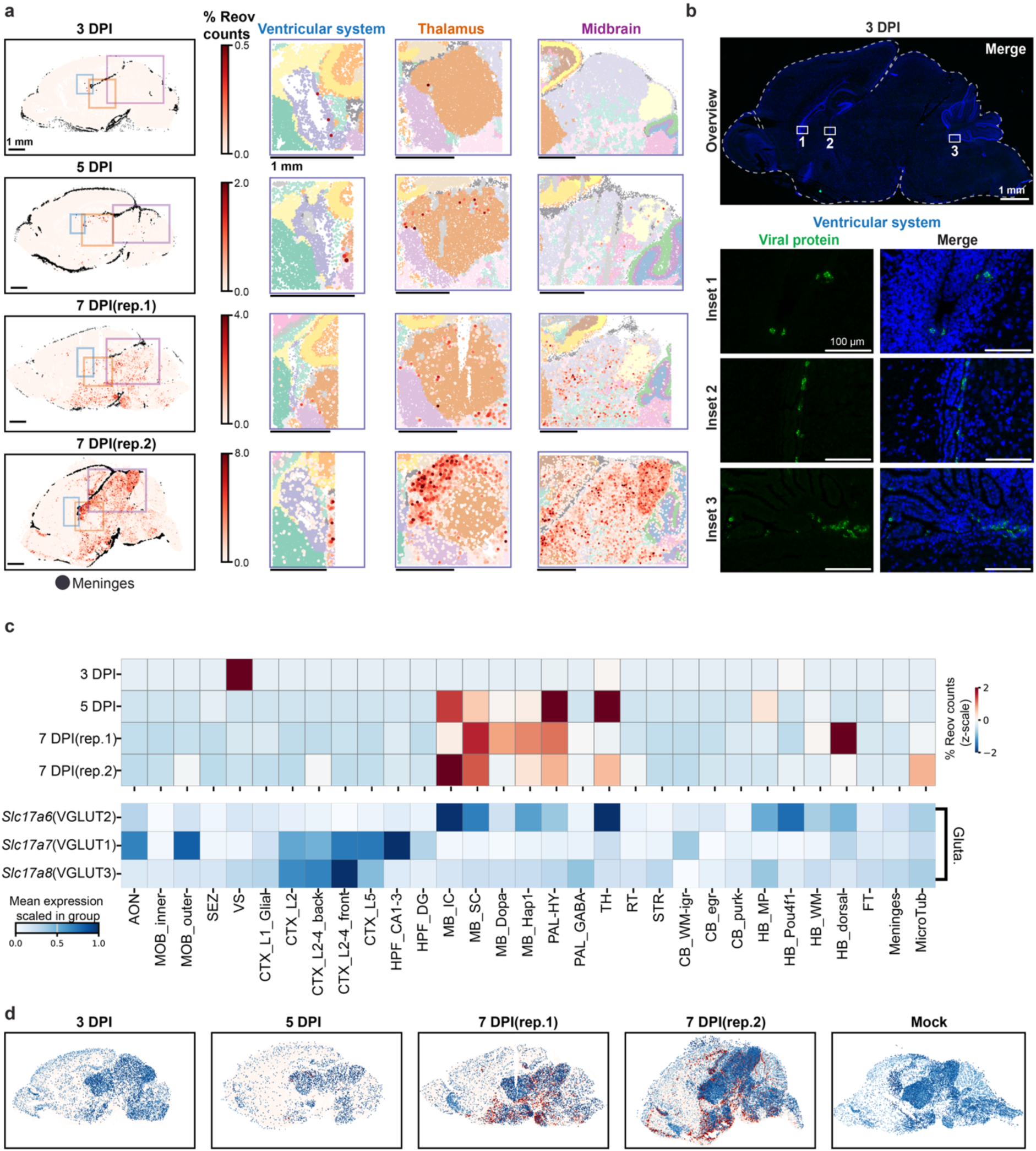
Spatiotemporal mapping of reovirus distribution reveals tropism progression in the neonatal mouse brain. **a** Spatial distribution of reovirus transcripts during the infection time course. Left panels display whole-brain maps of reovirus transcript abundance (% Reov counts relative to total counts). Colored boxes indicate regions of interest, ventricular system (blue), thalamus (orange), and midbrain (purple). Right panels show enlarged views of these regions with anatomical subregions overlaid. Scale bars, 1 mm. **b** Viral protein detection (green) in the ventricular system of a 3 DPI brain using indirect immunofluorescence (nuclei, blue). Top panel displays a representative whole-brain section (scale bar, 1 mm); bottom panels indicate zoom insets to the lateral (1), lateral-to-third (2), and fourth ventricle (3) regions (scale bars, 100 µM). **c** Heatmap comparing reovirus transcripts with glutamate neurotransmitter subtype transcripts across anatomical subregions. Top panel, z-scale percentage reovirus counts per subregion across infection time points. Bottom panel, mean expression levels of different vesicular glutamate transporters *Slc17a7* (VGLUT1), *Slc17a6* (VGLUT2), and *Slc17a8* (VGLUT3). **d** Spatial distribution of reovirus transcripts abundance overlaid with *Slc17a6* (VGLUT2) abundance.

Reovirus T3SA+ targets neurons in the CNS^12^. At 5 and 7 DPI, we observed high reovirus loads in subcortical regions, particularly the thalamus and midbrain (**Fig. 2a,c**). To identify the neuronal populations enriched in these regions, we examined expression patterns of markers for major neuron classes: excitatory glutamatergic neurons (*Slc17a6*, *Slc17a7*, *Slc17a8*), inhibitory GABAergic neurons (*Gad1*, *Gad2*, *Slc32a1*), cholinergic neurons (*Slc18a3*), dopaminergic neurons (*Slc6a3*), and noradrenergic neurons (*Slc6a2*) (**Supplementary Fig. 2b**). This analysis revealed that these infected regions are enriched for excitatory glutamatergic neurons. To determine whether a specific excitatory neuron subtype dictates viral tropism, we compared reovirus distribution with that of distinct glutamatergic subpopulations, as defined by vesicular glutamate transporter (VGLUT) genes^25^. In the mouse brain, VGLUT1 (*Slc17a7*) is the predominant transporter in cortical projection neurons, VGLUT2 (*Slc17a6*) characterizes subcortical glutamatergic populations, and VGLUT3 (*Slc17a8*) marks specific subsets of interneurons^25,26^. Cortical regions, which express high levels of *Slc17a7* (VGLUT1), had low viral loads throughout the infection time course. *Slc17a8* (VGLUT3) expression was sparse across most brain regions and did not correlate with viral distribution. In contrast, the spatial distribution of reovirus transcripts aligned with *Slc17a6* (VGLUT2) expression (**Fig. 2c**). Subregions with high reovirus loads at 5 and 7 DPI, including the thalamus, midbrain, and hindbrain, had high levels of *Slc17a6* (VGLUT2) expression (**Fig. 2c**). Visualization of reovirus transcripts combined with *Slc17a6* transcripts further support the overlapping distribution of viral infection with this neurotransmitter transcript (**Fig. 2d**). This suggests that reovirus preferentially targets VGLUT2-expressing neuronal populations, indicating subtype-specific tropism within excitatory neurons.

### Virus-host co-expression network analysis identifies distinct host immune activities in response to reovirus

To define host responses to reovirus infection in the mouse brain, we conducted an unbiased network-based analysis to uncover spatially co-expressed gene modules. We used our previously established Smoothie computational pipeline^27^ (**Fig. 3a**) to build and cluster a gene co-expression network for the 7 DPI replicate 2 brain, which had the highest density of viral RNA. The complete atlas consists of 3,586 genes divided into 121 modules of two or more genes (**Fig. 3b**). The atlas of all spatial gene modules is plotted in **Supplementary Figure 3**, and gene module members are listed in **Supplementary Data 1**. Genes within a given module have similar spatial patterning and frequently correspond to tissue regions, cell types, gene programs, or other conditions.

**Fig. 3.**
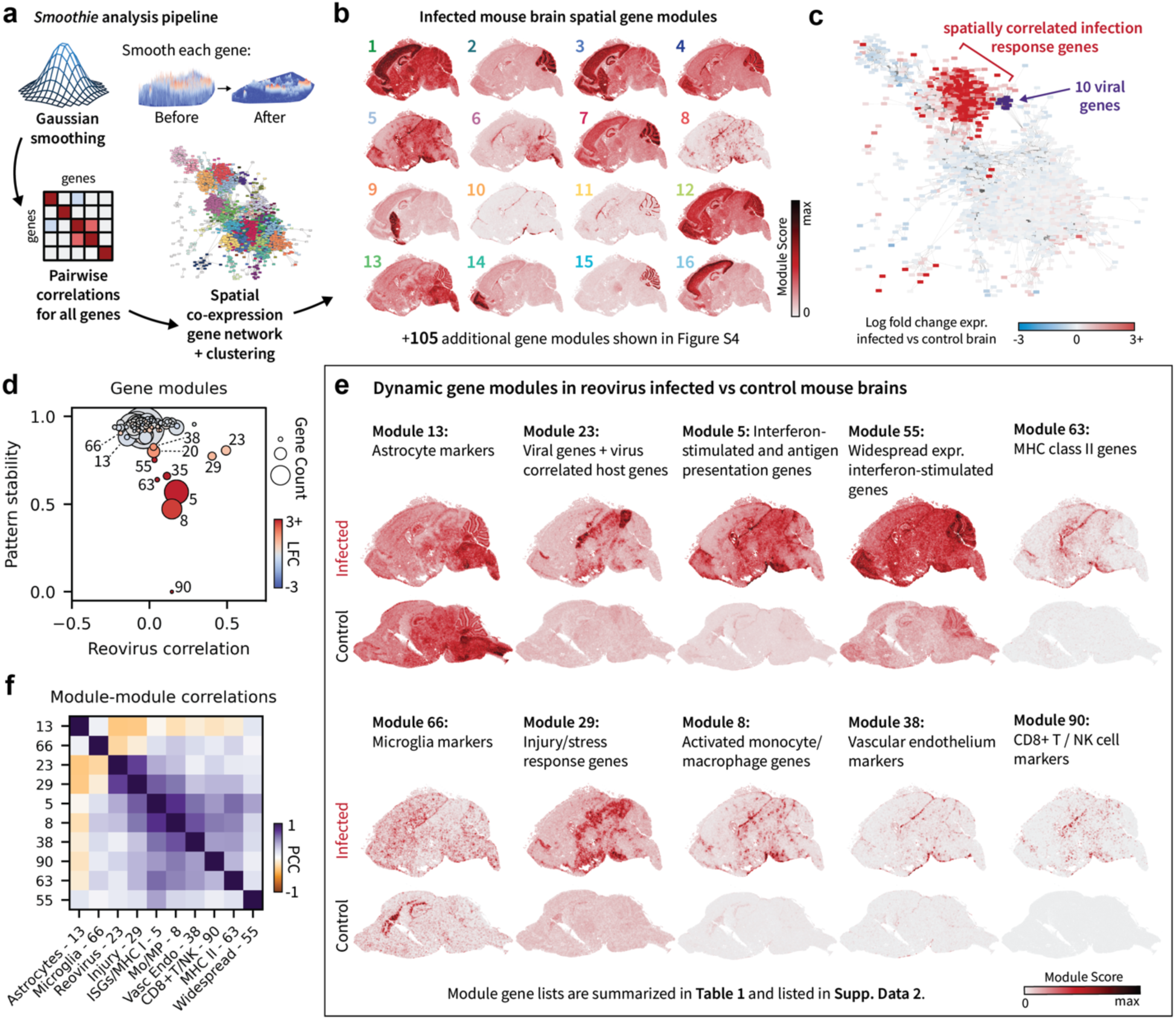
Reovirus-host spatial gene co-expression analysis identifies dynamic host gene modules following viral infection in the mouse brain. **a** Overview of the *Smoothie* analysis pipeline conducted using the day 7 post infection (dpi) mouse brain (replicate 2). Gaussian smoothing is used for spatial data denoising. Pairwise Pearson R correlation between gene expression levels provides a network for gene module clustering. **b** Example spatial expression visualizations of the 16 largest gene count modules of 121 total modules. All modules are shown in **Supplementary Fig. S3**. **c** Visualization of expression level bulk log*_e_* fold change (LFC) in the infected 7 dpi brain vs. the control brain reveals co-expression among many upregulated genes adjacent to the 10 reovirus RNAs. (Log base *e*). **d** For each of the 121 gene modules, median values of (i) transcript level correlation with reovirus infection, (ii) pattern stability, and (iii) LFC identify the gene modules with spatially distinct or dynamic patterns in infected vs. control brains. Color and size of points reflect bulk LFC (base *e*) and module gene count, respectively. **e** Visualization of dynamic host gene modules demonstrate differences between infected and control brains. Module annotations were inferred from gene set enrichment analysis using Enrichr. **f** Pairwise Pearson correlation coefficient (PCC) between module scores reveals correlation among many modules and anti-correlation of module 23 (containing reovirus RNAs) with modules 13 and 66 (astrocyte and microglia markers, respectively.) **b** and **e**, Module scores represent transcript averages after rescaling the transcript levels of each gene maximum value to 1.

To examine which of these 121 spatial gene modules are linked to a reovirus-specific host response, we projected the bulk log*_e_* fold change (LFC) of the transcript count of each gene onto the gene co-expression network. This analysis showed that many host genes upregulated in the infected brain are neighbors in the gene co-expression network closest to the 10 reovirus genes (**Fig. 3c**), indicating that most transcriptional changes occur near sites of reovirus infection. Apart from LFC, we also calculated the spatial correlation of each host gene’s RNA with reovirus RNA, and we measured a gene pattern stability metric that quantifies relative changes in the spatial distribution of each gene between the infected and control samples using second-order correlations. Using these three metrics – LFC, reovirus correlation, and pattern stability – we determined which of the 121 host gene modules differ between infected and control samples. Specifically, we plotted median gene values for each module across the three metrics and calculated robust Mahalanobis distances to identify statistically significant outlier modules (**Fig. 3d**). From this analysis, we highlight 10 modules as dynamic **(Fig. 3e)**, as these modules show clear spatial differences between the infected and control samples. To discover the underlying functions of the dynamic gene modules, we used Enrichr^28^ to compare the modules against several large databases for gene pathways, gene ontology, and cell types^29–33^. We also assembled a comprehensive table of highlighted gene families and pathways for each dynamic gene module (**Table 1**). In all, this approach allowed us to identify and annotate the dynamic gene modules between the infected and control samples, revealing a range of different spatially defined immune response gene programs that we discuss in the next paragraph.

**Table 1.**
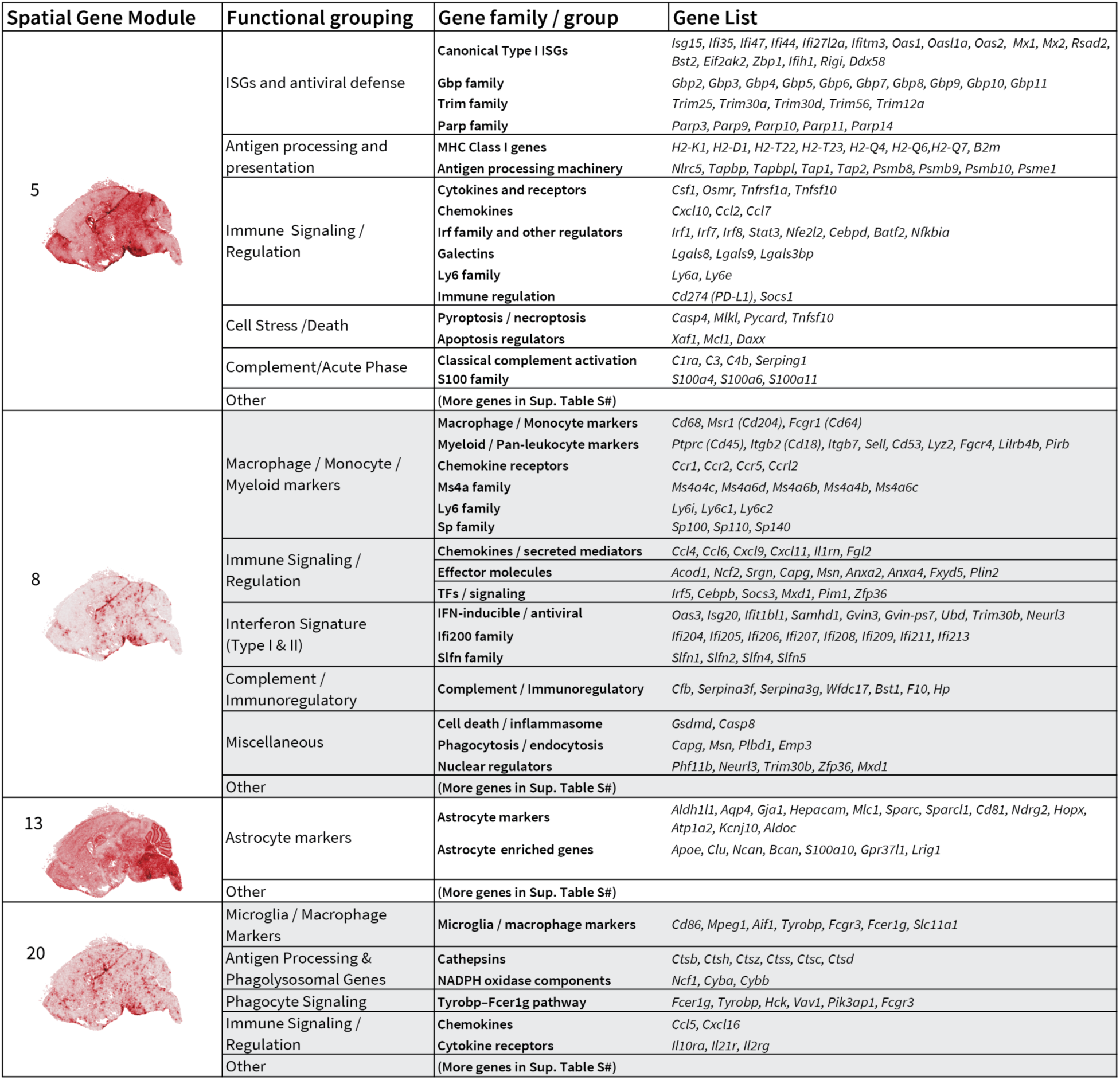
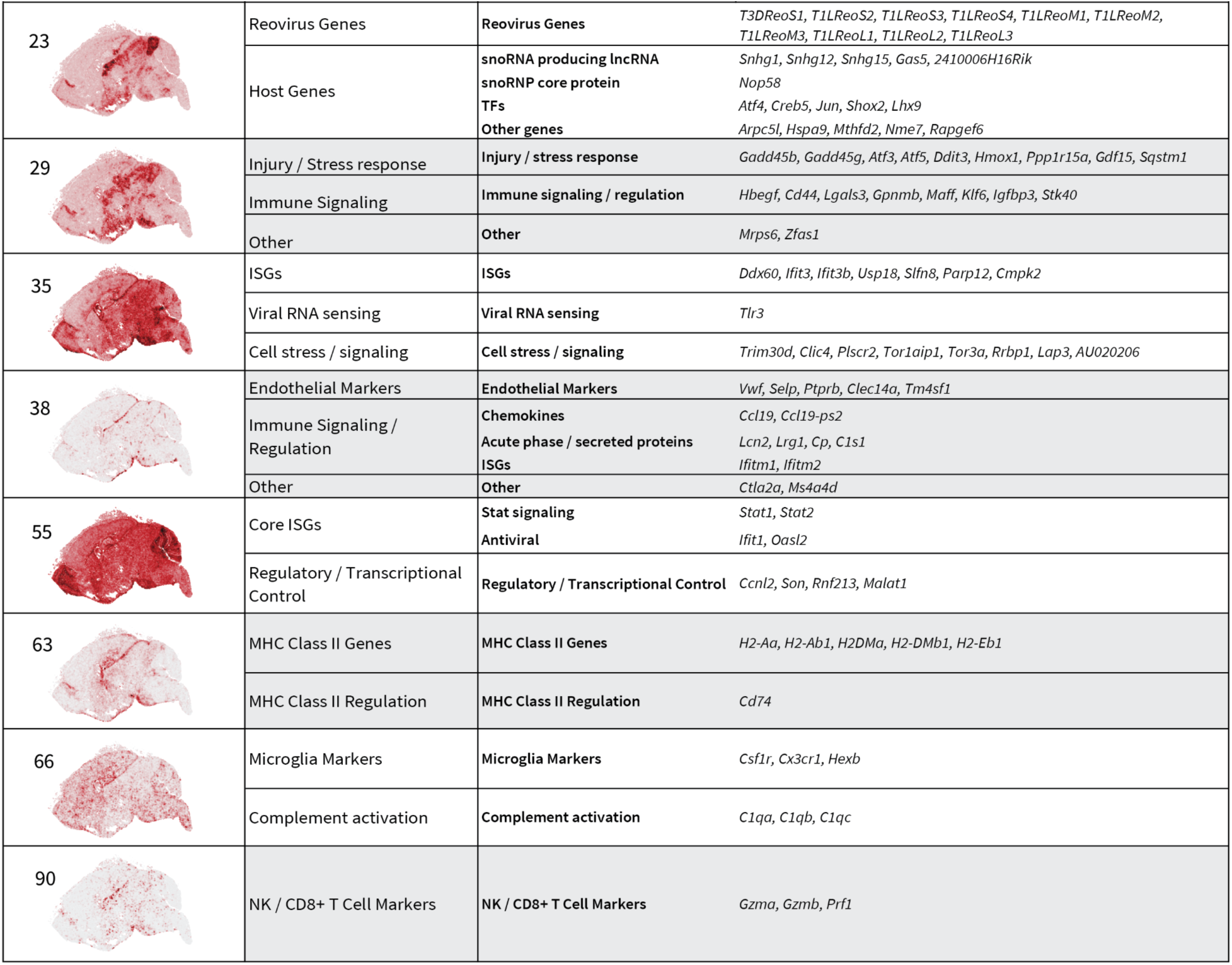
Gene lists and annotations for the spatial gene modules of dynamic host genes upon reovirus infection. Spatial gene modules with differences in pattern between control and infected samples. Dynamic modules are identified from the 121 total spatial modules by plotting median gene values for each module across three metrics, LFC, reovirus correlation, and pattern stability, and identifying outlier modules.

We found that dynamic gene modules between the infected and control samples corresponded to different aspects of the host immune response (**Table 1)**. For example, module 5 (176 genes) includes mostly antiviral and canonical type I interferon-stimulated genes (ISGs), including immune signaling genes (*PD-L1, Socs1, Stat3*), antigen-processing genes (MHC class I, *Psmb* family, *Tap* family), and other overlapping gene families (galectins, *Gbp* family, *Parp* family, *Trim* family). Module 8 (122 genes) contains marker genes for myeloid lineage cell types, particularly monocytes and macrophages (*Ccr2, Cd68, Fcar1, Msr1*), and a range of immune signaling, transcription factor, secreted molecule, and receptor genes (*Cebpb, Ccl6, Ccl4, Ccr1, Ccr2, Ccr5, Ccrl2, Cxc19, Cxcl11, Fgl2, Il1rn, Irf5, Socs3,*). Module 8 also includes several members of the *Ifi200, Ly6, Ms4a, Slfn, and Sp* gene families. Other dynamic modules include module 13 (95 genes, astrocyte markers), module 20 (46 genes, microglia and macrophage markers), module 23 (all 10 reovirus genes and 16 correlated host genes), module 29 (20 genes, cell injury and stress), module 35 (16 genes, ISGs), module 38 (15 genes, vascular and endothelial), module 55 (8 genes, ISGs), module 63 (6 genes, MHC Class II genes), module 66 (5 genes, microglia), and module 90 (3 genes, cytotoxic T cell and NK cell markers). Collectively, these findings reveal a coordinated immune response network during late-stage reovirus encephalitis.

Finally, we assessed module-by-module correlations in the 7 DPI replicate 2 sample to determine how reovirus-dependent modules spatially interact (**Fig. 3f**). This analysis revealed that module 13 (astrocyte markers) and module 66 (microglia markers) were anti-correlated with module 23 (reovirus transcripts), suggesting that glial cells are transcriptionally dysfunctional or nonviable in infected tissue. Module 29 (cell injury and stress) strongly correlated with module 23 (reovirus and host transcripts). Module 8 (macrophage and monocyte markers) correlated with module 38 (vascular endothelial markers). These correlations and anti-correlations also are visually striking in the spatial plots (**Fig. 3e**). Together, these analyses comprehensively identify dynamic patterns of host gene expression during reovirus infection and cluster infection-associated genes into precise spatial modules that capture different facets of the host immune response.

### Several snoRNA-related transcripts strongly correlate with reovirus, while astrocyte markers exhibit viral anti-correlation

Sixteen host genes clustered with reovirus RNAs in gene module 23 (**Table 1**). Of these 16 genes, five were transcription factors (*Atf4, Creb5, Jun, Lhx9, Shox2*), five have metabolic, signaling, or structural functions (*Arpc5l, Hspa9, Mthfd2, Nme7*, *Rapgef6*), and six were linked to small nucleolar RNA (snoRNA) biogenesis (*Gas5, Nop58, Snhg1, Snhg12, Snhg15, 2410006H16Rik*). The spatial correlation between reovirus and snoRNA biogenesis genes is interesting, as host snoRNAs influence replication of other RNA viruses, including dengue virus, human rhinovirus 16, and influenza virus^34^. Visualizing four example snoRNA biogenesis transcripts (*Snhg1, Snhg12, Snhg15, 2410006H16Rik*), there is a striking resemblance to the spatial distribution of reovirus transcripts for both day 7 infected replicates (**Fig. 4a**). In contrast, when examining the day 7 control sample, these snoRNA gene patterns are absent. Rank plots of top reovirus-correlated genes show that *Gas5, Nop58, Snhg1, Snhg12, Snhg15, Snhg17,* and *2410006H16Rik* were highly correlated with reovirus gene expression (**Fig. 4b**).

**Fig. 4.**
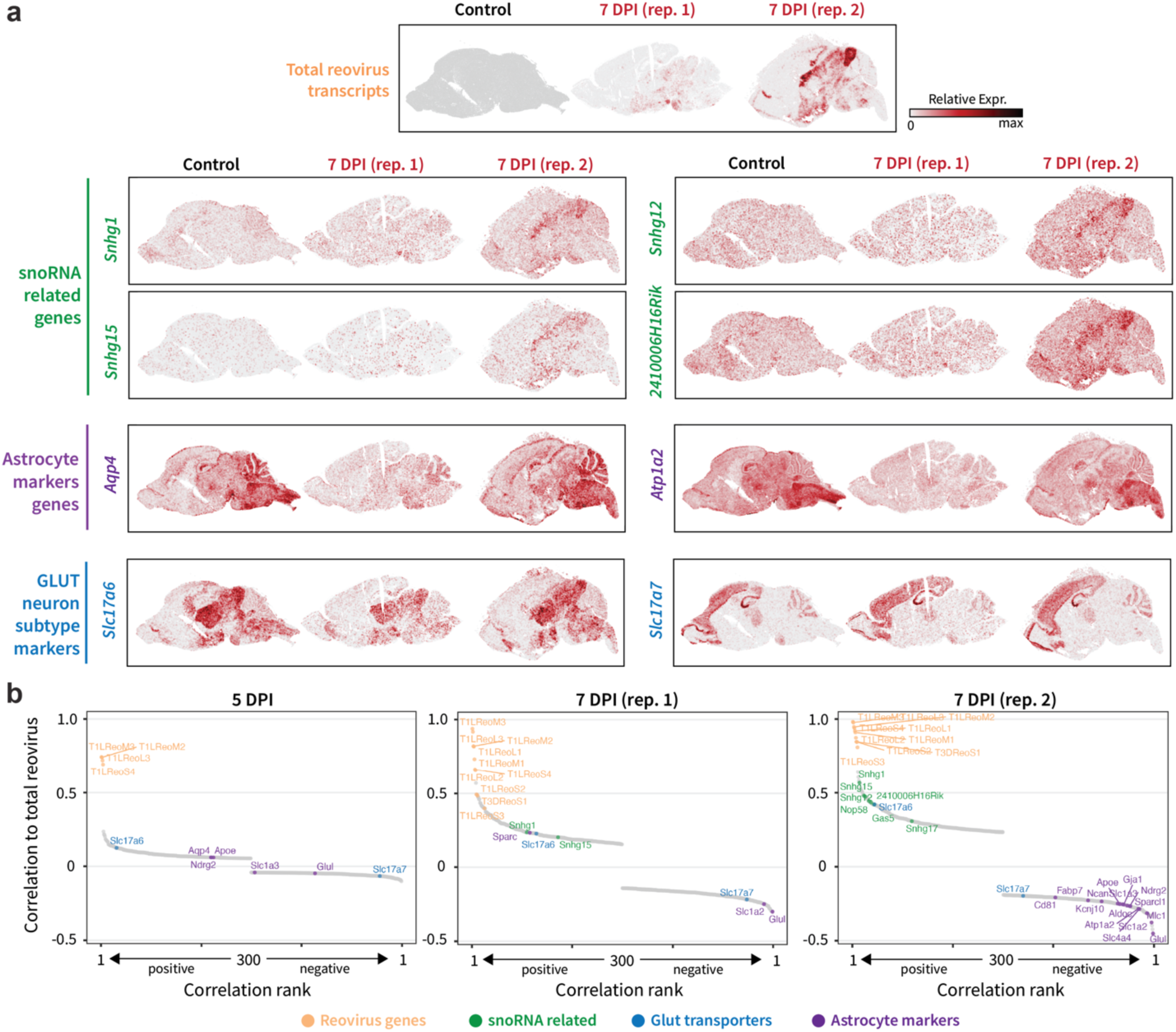
Expression of host snoRNA-related genes positively correlates with reovirus RNAs, while expression of astrocyte marker genes is anti-correlated. **a** Spatial expression plots of total virus transcripts, snoRNA-related transcripts (*Snhg1*, *Snhg12*, *Snhg15*, and *241006H16Rik*), astrocyte-marker transcripts (*Aqp4* and *Atp1a2*), and VGLUT transcripts (*Slc7a6* and *Slc7a7*). **b** Reovirus correlation and anti-correlation rank plots show positive correlation with expression of snoRNA-related genes and anti-correlation with expression of astrocyte-marker genes.

Meanwhile, astrocyte-enriched gene module 13 showed a negative correlation with the spatial distribution of reovirus transcripts (**Table 1**). Spatial maps of canonical astrocyte marker genes *Aqp4* and *Atp1a2* revealed clear patterns that were anti-correlated with reovirus transcripts (**Fig. 4a**). Correlation rank plots consistently demonstrated negative associations between reovirus transcripts and several additional astrocyte-enriched genes in module 13, including *Aldh1l1, Aldoc, Aqp4, Atp1a2, Gja1, Glul, Hepacam, Mlc1, Ndrg2, Sparc,* and *Sparcl1)* (**Fig. 4b**). This finding was unexpected, as reovirus is thought to predominantly infect neurons rather than astrocytes. The inverse spatial relationship between viral transcripts and astrocyte marker expression indicates that astrocyte gene expression is suppressed in reovirus-infected areas. This pattern may reflect local astrocyte dysfunction, transcriptional silencing, or astrocyte loss, presumably because of viral replication, in infected areas. Finally, consistent with earlier observations (**Fig. 2c,d**), the spatial distribution of transporter gene *Slc7a6* (VGLUT2) transcripts showed a positive correlation with viral RNA, demonstrating preferential viral localization within excitatory neuronal populations (**Fig. 4a,b**).

### Spatiotemporal development of the host infection response in the brain

To understand how the host response to reovirus infection developed over time, we first calculated an infection-response score for each of the 3-, 5-, and 7-day infected samples relative to the control sample. The mean infection response score per anatomical subregion showed that at 3 and 5 DPI, responses were mostly localized to the vasculature and meninges, but by 7 DPI, responses extended into the brain parenchyma (**Fig. 5a**). Next, for each infected sample we identified spatial submodules of genes within the overall infection response using graph-based clustering on gene co-expression networks. Gene modules with dynamic expression relative to the control were identified as described in **Figure 3c**, and the dynamic modules are highlighted in the gene co-expression networks in **Figure 5b**. Over the course of infection, the number of dynamic modules increased markedly (**Fig. 5c**). At 3 DPI, we detected only one large dynamic gene module that was composed primarily of ISGs in the vasculature and meninges (**Fig. 5d**). At 5-7 DPI, this large ISG module persisted, and new spatial gene modules appeared. At 5 DPI, a module corresponding to reovirus genes and a meninges-associated module containing macrophage and myeloid markers was evident (**Fig. 5c,d**). By 7 DPI, additional modules emerged that corresponded to macrophage and microglia marker genes, injury and stress genes, MHC class II genes, and markers of NK cells and CD8+ T cells (**Fig. 5c,d**). In 7 DPI samples, expression of the large ISG module expanded across the brain, while the newly emerging modules were localized to areas of viral infection (**Fig. 5d)**. Examining changes in bulk expression across time points further illustrates the temporal activation of immune gene modules. We plotted the median scaled gene expression value for modules 5, 8, 55, 63, and 90 across datasets (**Fig. 5e**), which revealed early and late signatures of reovirus encephalitis. The individual scaled gene expression values from **Figure 5e** are provided in **Supplementary Fig. 4**. Taken together, these findings allow us to order the reovirus-induced host immune response from (i) elevated ISG expression to (ii) macrophage and myeloid infiltration to (iii) MHC class II expression and cytotoxic lymphocyte infiltration.

**Fig. 5.**
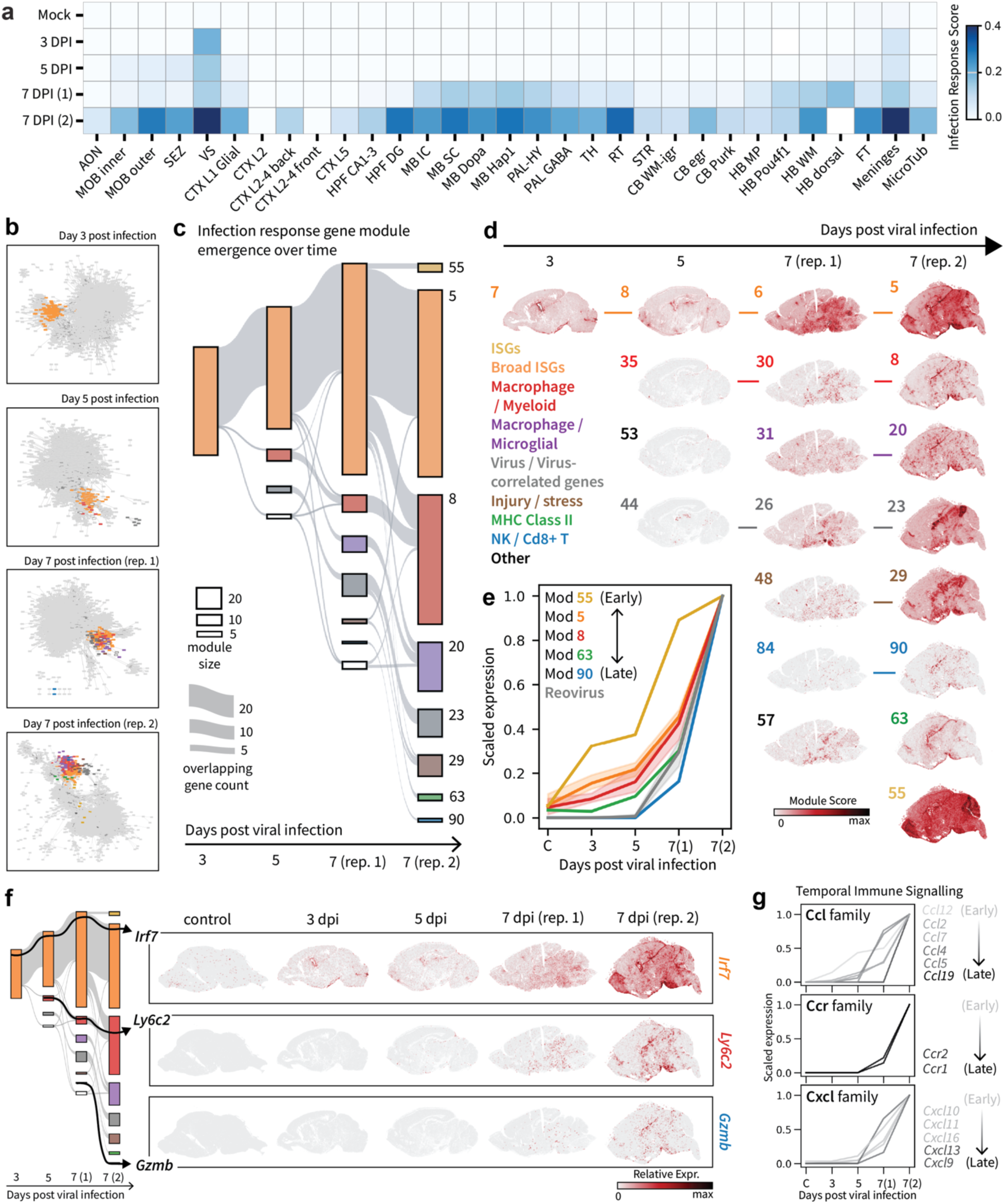
Spatiotemporal host immune response over the course of reovirus infection in the neonatal mouse brain. **a** Heatmap of mean infection response scores across brain anatomical subregions across samples. The infection response score is an average of the top upregulated genes for each infected sample compared with control. **b** Virus-host correlation networks produced for each individual infected brain sample. Colorful modules are dynamic modules that show different expression levels relative to the control brain. **c** Sankey plot showing the overlap and emergence of dynamic gene modules as reovirus infection progresses, highlighting the cadence of more distinct gene modules. Rectangle height and connecting line height reflect the size of the gene modules and gene module overlaps, respectively. **d** Spatial gene module plots of dynamic gene modules for each infected sample. Labels mark each temporal gene module based on the final 7 DPI (high) gene module sets. **e** Temporal comparison of relative bulk expression for each of the highlighted 7 DPI (high) dynamic gene modules. Lines represent median values within the gene set, and 95% CIs are shown for the medians for groups with > 10 genes. **f** Example spatial plots for three dynamic genes extracted from the Sankey plot. Interferon-stimulated gene *Irf7* is a dynamic gene starting from 3 DPI. Macrophage marker gene *Ly6c2* is a dynamic gene starting from 5 DPI. NK and CD8+ T cell marker *Gzmb* is a dynamic gene in the 7 DPI infected samples. **g** Temporal expression ordering with bulk relative expression plots for chemokines and the Ccr receptor family. Some chemokine genes, such as *Ccl12*, are expressed as early as 3 DPI, whereas others like *Ccl19*, are expressed at 7 DPI.

By spatiotemporally ordering these gene modules, we were able to examine selected genes of interest to virus-host interactions. For example, we observed that the master immune regulator *Irf7* is expressed from 3 DPI onward, the macrophage marker *Ly6c2* is expressed from 5 DPI onward, and the apoptosis-initiating enzyme granzyme B (*Gzmb*) is expressed at 7 DPI (**Fig. 5f**). The complete list of dynamic genes from **Figure 5c** and their order of emergence is listed in **Supplementary Data 2**. We also examined expression trajectories for gene families of interest. To illustrate this approach, we determined the temporal expression trajectories of the *Ccl* and *Cxcl* chemokine families and *Ccr* receptors, which regulate leukocyte recruitment during infection. *Ccl12* was induced first at 3 DPI, followed by *Ccl2* and *Ccl7* at 5 DPI, and *Ccl4, Ccl5*, and *Ccl19* at 7 DPI (**Fig. 5g**). Similarly, *Cxcl10, Cxcl11*, and *Cxcl16* were expressed from 5 DPI, while *Cxcl9* and *Cxcl13* emerged at 7 DPI alongside *Ccr1* and *Ccr2* (**Fig. 5g**). These patterns reveal a staged immune program in the infected brain, progressing from early CCR2-mediated monocyte recruitment (*Ccl12→Ccl2/7*) to interferon-driven T cell trafficking (*Cxcl10/11/16*) to an additional wave of adaptive immune recruitment at 7 DPI (*Ccl4/5/19, Cxcl9/13*). While we highlight immune signaling molecules here, this temporal bulk expression analysis can be conducted for any gene family in the datasets, and we extend the analysis to 63 gene name prefixes (families) within the dynamic infection response modules in **Supplemental Fig. 6**. In all, by spatiotemporally ordering host immune response genes, we provide a rich resource to examine genes or gene families of interest during a time course of viral encephalitis.

## Discussion

In this study, we produced a high-resolution spatiotemporal transcriptomic atlas of reovirus encephalitis, capturing dynamic interactions between viral propagation and host immunity in the neonatal mouse brain. Our spatial data suggest a defined route of viral dissemination. At the earliest time point examined (3 DPI), viral transcripts were confined to ventricular boundaries before spreading into the thalamus and midbrain by 5 and 7 DPI. This ventricle-to-parenchyma trajectory supports a model in which reovirus exploits cerebrospinal fluid (CSF) circulation to access the central nervous system. As the principal site of CSF production, the choroid plexus may serve as a natural entry point for reovirus into the CNS, enabling viral exposure to neuronal populations adjacent to the ventricular system. From these periventricular regions, viral burden expands preferentially within defined subcortical territories.

Regions of high viral load spatially colocalize with expression of *Slc17a6* (VGLUT2). Although both subcortical and cortical structures lie adjacent to ventricular zones and should, in principle, have comparable exposure to CSF-borne virus, reovirus preferentially accumulates in regions of VGLUT2⁺ neurons rather than VGLUT1⁺ populations. Cross-species transcriptomic atlases demonstrate that *Slc17a6* (VGLUT2) marks an evolutionarily conserved subcortical glutamatergic neuron population, distinct from the mammalian-specialized *Slc17a7* (VGLUT1) neocortical lineage^26^. This evolutionary divergence may correspond to differences in surface receptors, intracellular trafficking pathways, or antiviral defense programs that influence susceptibility to viral infection. Direct comparative studies of VGLUT1⁺ and VGLUT2⁺ neurons will be required to define the molecular basis of this subtype-specific tropism.

We used unbiased gene co-expression networks to resolve the host immune response into spatially defined modules. This analysis enabled us to construct a detailed table of gene sets per module, which serves two purposes, (i) annotating the spatial modules with marker genes and (ii) highlighting new gene family associations within the modules. Further analysis revealed a chronological layering of immune activity. The response is first detected in our study at 3 DPI with a module of ISGs, e.g., *Irf7* and *Stat3*, centered on the vascular and meningeal interfaces. This early response is followed at 5 DPI by the emergence of myeloid-specific modules, e.g., *Ccr2*, *Cd68*, and *Ly6c2*, likely reflecting infiltration of monocytes from the bloodstream and an expansion or redistribution of macrophages from the brain. By 7 DPI, which is a peak time of disease, the immune landscape becomes more complex, featuring the appearance of cytotoxic T-cell and NK cell signatures, e.g., *Gzma*, *Gzmb*, and *Prf1*, marking a shift to cell killing and adaptive immunity. The temporal activation of chemokine gene expression mirrors these findings, progressing from early CCR2-mediated monocyte recruitment (*Ccl12* at 3 DPI, *Ccl2/7* at 5 DPI) to interferon-driven T cell trafficking (*Cxcl10/11/16* at 5 DPI) to additional adaptive immune recruitment signals (*Ccl4/5/19, Cxcl9/13* at 7 DPI). These results reveal the regulation of the antiviral response, evolving from innate alarm signals to targeted cellular cytotoxicity.

Perhaps the most striking finding from our network analysis is the strong positive correlation between reovirus replication and host small nucleolar RNA (snoRNA) biogenesis genes (*Gas5*, *Nop58*, *Snhg1*, *Snhg12*, *Snhg15*, *Snhg17*, and *2410006H16Rik*). snoRNAs are non-coding RNAs that form complexes with proteins (snoRNPs) to guide the chemical modification (e.g., methylation, pseudouridylation) of ribosomal RNAs (rRNAs). While other viruses use snoRNA machinery, there are limited reports of the use of this system by viruses containing double-stranded RNA genomes, such as reovirus. Notably, a prior gene-trap screen showed that disruption of the snoRNA host gene SNHG2 confers resistance to reovirus^35^. In our study, we identify seven additional snoRNA-associated genes with spatial expression patterns that suggest a potential role in reovirus infection. One possibility is that reovirus co-opts host ribosomal modification machinery to optimize translation of viral mRNAs or to stabilize viral mRNA secondary structures. These observations nominate snoRNA-associated pathways as previously unrecognized components of reovirus–host interactions and warrant further investigation.

Not all host gene expression programs positively correlate with viral burden. We observed a strong inverse relationship between reovirus transcripts and astrocyte markers, including *Aldh1l1*, *Aqp4*, and *Mlc1*. This pattern may reflect intrinsic resistance of astrocytes to reovirus infection^36^, leading to viral exclusion, or alternatively, localized astrocyte dysfunction or loss in regions of high viral replication. Downregulation of genes such as *Mlc1*, which contributes to blood-brain barrier integrity and ion homeostasis, suggests that viral replication may disrupt astrocytic end-feet and glutamate clearance. Such perturbations could exacerbate excitotoxicity and contribute to seizures observed in severe viral encephalitis^37^.

Collectively, our findings enable construction of a molecularly resolved map of reovirus encephalitis. By delineating spatially restricted viral tropism, defining neuronal subtype-specific localization of virus, and resolving the stepwise activation of innate and adaptive immune programs, this atlas provides a comprehensive framework for understanding how neurotropic viruses navigate and disrupt the developing brain.

## Methods

### Ethical approval for animal experiments

All animal husbandry and experimental procedures were conducted in accordance with US Public Health Service policy and were approved by the Institutional Animal Care and Use Committees at Cornell University (IACUC no. 2019-0129) and the University of Pittsburgh (IACUC no. 25046563).

### Reovirus infection of neonatal C57BL/6J mice

Three-day-old (P3) mice of both sexes were inoculated intracranially into the right hemisphere with 300 plaque-forming units (PFU) of reovirus strain T3SA+ diluted in phosphate-buffered saline (PBS). Independent litters were inoculated with an equal volume of PBS by the same route and served as mock controls. Mice were monitored daily, and at 3, 5, or 7 days post-inoculation, animals were euthanized, and brain tissues were harvested aseptically. For each animal, the right hemisphere was processed for spatial transcriptomics and histological analyses, and the left hemisphere was used for quantifying viral titers by plaque assay^24^. Right brain hemispheres were embedded in OCT media (SAKURA, 25608-930) and flash-frozen in a liquid-nitrogen-cooled isopentane (EMD Millipore, MX0760) bath. Tissue blocks were sagittally sectioned using a cryostat.

### Slide-seq spatial transcriptomics sample preparation and library generation

Right brain tissue sections (10-µm thick) were mounted on 1 × 1 cm Curio Seeker tiles. A barcode whitelist and a barcode position file for the corresponding tile were provided by Curio Bioscience. Slide-seq spatial transcriptomics experiments were conducted using the Curio Seeker Kit according to manufacturer’s instructions. Libraries were sequenced using an Illumina NovaSeq 6000 instrument with paired-end 150 bp reads (PE150). The sequencing reads were aligned to a *Mus musculus* (mm39)^38^ genome and recombinant reovirus reference genome (T3SA+)^12^ using the slide-snake pipeline (GitHub: https://github.com/mckellardw/slide_snake) to derive a feature-by-bead barcode expression matrix.

### Slide-seq data pre-processing

The slide-seq count matrix and position information for every bead barcode were placed into an AnnData^39^ object using Scanpy^40^ after filtering the low-quality beads with fewer than 100 unique molecular identifier (UMI) counts. To reduce the smear effect inherent in Slide-seq capture, beads were aggregated into 40-um spatial bins by summing spots count within each grid cell. Bins with less than 5 spots inside the grid cell were removed. For each bin, we calculated pairwise Euclidean distances to all other bins using scipy.spatial.distance.pdist and enumerated spatial neighbors within a 100-unit radius. Bins with fewer than 5 spatial neighbors within the radius were excluded from subsequent analyses.

### Multilevel anatomical region annotations

A customized GraphST-based approach was used for anatomical region annotations to leverage gene expression and spatial proximity information. Spatial spots were aggregated into 30-µm bins by summing UMI counts across beads within each grid cell. All samples were combined, and highly variable genes (HVGs) were identified using the Seurat v3 method^41^ (n_top_genes=3000) with batch correction across samples. To prepare the data, we normalized counts (CPM) and log-transformed the data to stabilize variance. We then restricted the analysis to the most informative, highly variable genes (HVGs) and scaled the expression levels following GraphST recommendations. For each sample, spatial graphs were prepared using the Delaunay triangulation method^42^ with 50-µm radius pruning. GraphST models were trained on each sample separately to yield sample-specific spatial embedding layers (obsm[‘emb’]) that encode both transcriptomic and spatial neighborhood information. Principal component analysis (PCA, n_components=25) was used on the concatenated embeddings from all samples. Harmony integration^43^ was conducted using the PC space to correct for batch effects between samples with the sample identity as the batch key. A k-nearest neighbor graph was prepared using the Harmony-integrated PCs (n_pcs=20), and Leiden clustering^44^ was conducted to identify anatomical clusters (resolution=1.4). Differential expression analysis (Wilcoxon rank-sum test) was used to identify top marker genes for each cluster. Clusters were manually annotated based on spatial localization patterns and marker gene expression, cross-referencing with brain anatomy and the Allen Mouse Brain Common Coordinate Framework version 3 (CCFv3)^21,45^. This approach yielded 32 distinct anatomical regions in the brain with one cluster consisting of inflammatory response markers. Spots in this cluster were annotated as “Region_Inflam” and were further assigned to appropriate anatomical contexts based on spatial neighbor identity. Region labels from the binned analysis were mapped back to the individual beads using barcode matching for subsequent visualization and analysis.

### Multilevel cell type annotations

Cell type annotation followed a two-stage approach using deeply sequenced samples as internal spatial references. First, the Mock and 7 DPI (rep. 2) samples were combined and integrated using Harmony^46^ to correct for batch effects between samples. Leiden clustering was conducted (resolution = 1.8), and clusters were manually annotated by cross-referencing marker genes with established cell signatures from the Allen Mouse Brain Atlas^21^. This analysis identified 35 distinct cell types, including neuronal subtypes (glutamatergic, GABAergic, cholinergic, and dopaminergic neurons), glial cells (astrocytes, oligodendrocytes, and microglia), vascular cells, choroid plexus epithelial cells, and other brain-resident populations (**Supplementary Fig. 1d,e**). Second, the 3 DPI, 5 DPI, and 7 DPI (rep. 1) samples were assigned using PrismST deconvolution^22^ with the annotated Mock and 7 DPI (rep. 2) datasets serving as the reference. Prior to deconvolution, reference gene expression matrices were filtered using the cleanup.genes function to remove ribosomal protein genes, mitochondrial ribosomal protein genes, and sex chromosome-linked genes. Differentially expressed marker genes were identified using get.exp.stat with a minimum cell count cutoff of 20 cells per type, and markers were selected using select.marker with thresholds of *P* value < 0.05 and log fold-change > 0.01. Deconvolution was accomplished using run.prismST to estimate cell-type proportions (theta) for each spatial spot. Each spot was assigned to the cell type with the maximum estimated proportion. This spatial-to-spatial deconvolution strategy, rather than using external single-cell RNA-seq references, preserves platform-specific expression signatures and enables robust cell-type assignment across samples with varying sequencing depths.

### Indirect immunofluorescence for viral proteins

Right brain tissue sections (7-10-µm thick) were mounted on histology slides at −12 to −18°C and stored at −80°C until staining. Tissue sections were stained by indirect immunofluorescence to detect the reovirus σNS protein, which marks viral replication factories in infected cells. All subsequent incubations were conducted at room temperature, unless indicated otherwise. Slides were briefly thawed/dried before submersion in 4% paraformaldehyde. Tissue was permeabilized with PBS supplemented to contain 0.1% Triton X-100, washed with PBS, and blocked with PBS supplemented to contain 10% goat serum. Tissue was incubated at 4°C overnight with custom guinea pig antisera raised against purified reovirus σNS protein (1:3,000 in PBS supplemented to contain 1% bovine serum albumin). Tissue was washed 3 times with PBS and incubated with Alexa Fluor™ 488-conjugated goat anti-guinea pig secondary antibody for 1 h. Tissue was counterstained with DAPI, washed extensively, and mounted with coverslips. Individual fluorescent images were tile-scanned at 4× magnification (whole-brain overview) or 10× magnification (insets) using a Lionheart FX imager (Agilent Technologies). Automated image stitching was accomplished using Gen5 software (Agilent Technologies). FIJI software^47^ was used to prepare the final images.

### Smoothie gene co-expression network analysis for the 7 DPI (rep. 2) sample

Quality control for this analysis was accomplished by removing transcriptomes with less than 50 UMIs and genes with less than 100 UMIs. The data were normalized with counts per thousand (CPT) and log1p normalization. Gaussian smoothing was conducted with parameters *grid_based_or_not=False*, *gaussian_sd=30 microns*, and *min_spots_under_gaussian=25*. Gene network construction and clustering was accomplished using parameters *pcc_cutoff = 0.4* and *clustering_power = 10*.^27^

### Time course infection response score calculation

Infection response scores were identified using gene co-expression network analysis conducted separately for each timepoint. Genes within identified infection modules were ranked by network degree (connectivity), and the top 100 highest-degree genes from each infection module, 3 DPI, 5 DPI, 7 DPI (rep. 1), 7 DPI (rep. 2), were selected for scoring. The intersection of all gene sets from the infected samples was used for the mock control infection response score. Infection response scores were computed using scanpy’s score_genes function with the controlling gene set size matched with the signature gene list length for background correction. Scores were calculated independently for each batch to account for timepoint-specific network topology while maintaining consistent methodology across samples.

### Time course Smoothie gene co-expression network analysis

Quality control for the 3 DPI, 5 DPI, and 7 DPI (rep. 1) datasets involved removing transcriptomes with less than 20 UMIs and genes with less than 50 UMIs. The data were normalized with counts per thousand (CPT) and log1p normalization. Gaussian smoothing was conducted with parameters *grid_based_or_not=False*, *gaussian_sd=30 microns*, and *min_spots_under_gaussian=25*. To normalize gene retention, gene networks for each dataset were constructed using a *pcc_cutoff* value that yielded a network with the same number of genes as the 7 DPI (rep. 2) dataset with *pcc_cutoff = 0.4*. Clustering was conducted using *clustering_power = 10* for all datasets.

### Identifying dynamic gene modules in infected vs. control samples

Gene modules in each of the infected samples with altered expression relative to control were classified as *dynamic* modules. Modules were classified as dynamic by applying a robust Mahalanobis distance-based outlier test to median values for each gene across three dimensions (metrics): (i) gene transcript log fold-change (LFC), (ii) a second-order correlation gene *pattern stability* metric from the Smoothie pipeline^27^, and (iii) correlation of levels of each transcript with total reovirus RNA. The gene pattern stability metric of Smoothie was used to assess whether the gene maintains similar correlation profiles among other genes in the control and infected datasets. The metrics were chosen based on the following criteria: (i) LFC identifies the genes with bulk expression upregulation, which was observed for many immune activated genes, (ii) gene pattern stability identifies genes with changing correlations with the majority of other genes, indicating genes with shifts in their spatial distributions following infection, and (iii) correlation with reovirus suggests that expression of a host gene is influenced by viral infection. For Figure 3, all three metrics were used in Mahalanobis distance calculation. For Figure 5, only the first two metrics, LFC and gene pattern stability, were used in Mahalanobis distance calculation, as earlier timepoints (3 DPI and 5 DPI) lacked robust reovirus transcript expression patterns, rendering the third metric less accurate.

### Normalized bulk gene expression trajectory analyses

Normalized bulk gene expression was used to account for differences in sequencing depth across datasets. For each gene, expression was calculated as the fraction of total UMIs in each dataset, and these values were normalized by dividing by the maximum observed value across datasets, yielding gene-level values scaled to a maximum of 1.

## Data Availability

All raw sequencing data, spatial barcodes lists, and raw count matrices will be made publicly available upon publication.

## Code Availability

All scripts and code will be made publicly available on GitHub upon publication.

## Acknowledgements

We are grateful to members of the De Vlaminck, Dermody, and Parker laboratories for thoughtful discussions and suggestions. This study was supported by U.S. Public Health Service awards T32 AI049820 (K.A.G.), R01 AI176681 (J.S.L.P. and I.D.V.), R01 AI174526 (D.M.S., and T.S.D.), and NSF DGE 2139899 (C.H.). Additional support was provided by UPMC Children’s Hospital of Pittsburgh and the Heinz Endowments. The funders had no role in study design, data collection and interpretation, or the decision to submit the work for publication.

## Author Contributions

C.H. and S.J. conceived and designed experiments, conducted experiments, analyzed data, contributed materials/analysis tools, and wrote the paper. K.A.G., M.E.F., and K.M.K. conceived and designed experiments, conducted experiments, and analyzed data. P.S., M.M., and M.H. conceived and designed experiments and conducted experiments. D.M.S., J.S.L.P., T.S.D, and I.D.V. conceived and designed experiments, analyzed data, and wrote the paper. All authors reviewed, critiqued, and provided comments on the manuscript.

