## Supplementary Figures for "High-resolution spatial transcriptomics maps viral tropism and reveals spatially organized immune modules in viral encephalitis"

**Affiliations**

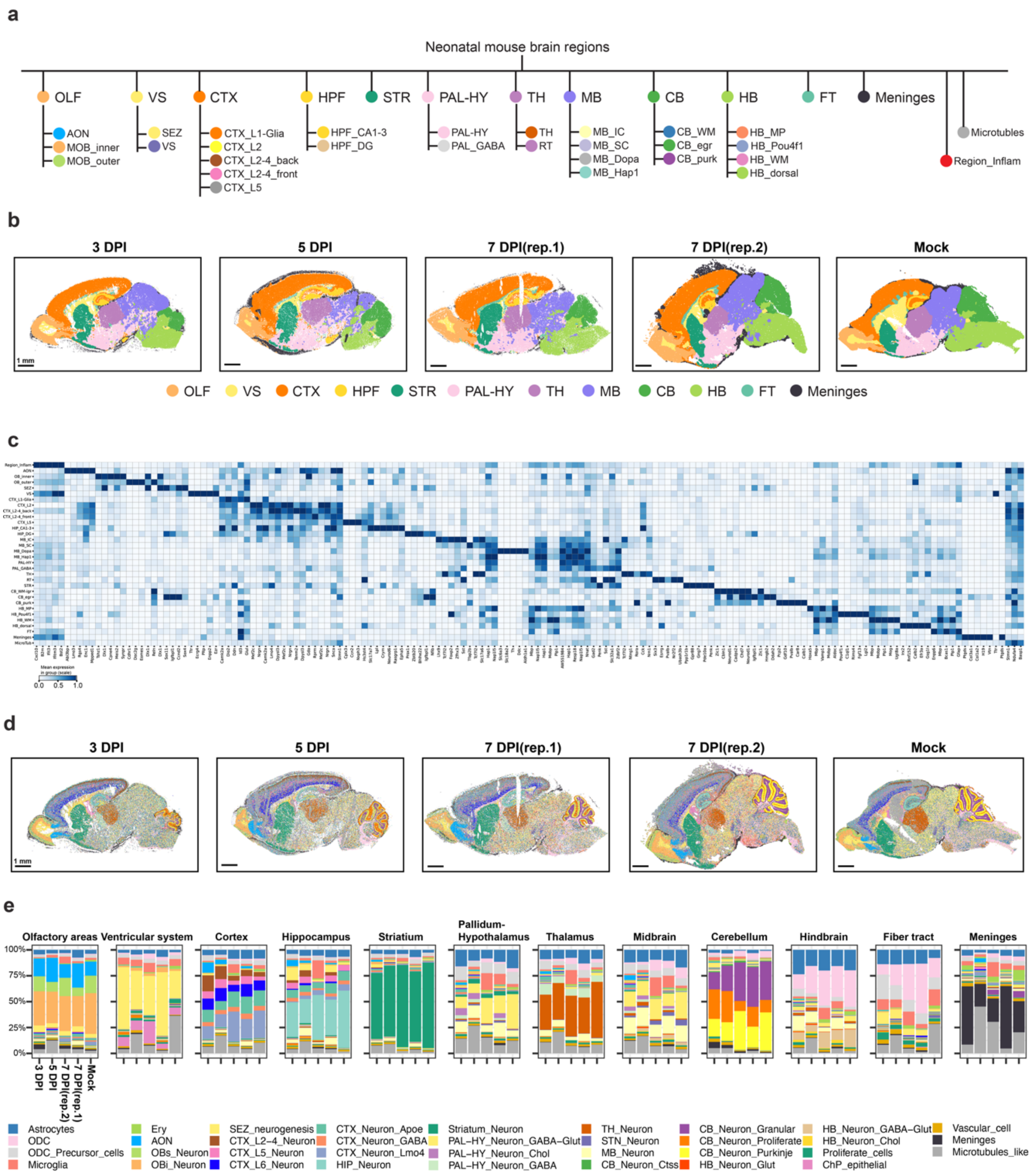


**Supplementary Fig. 1. a** Hierarchical organization of brain region annotation showing major anatomical regions (top level) and their constituent anatomical subregions (bottom level). OLF, olfactory; VS, ventricular system; CTX, cortex; HPF, hippocampus formation; STR, striatum; PAL-HY, pallidum-hypothalamus; TH, thalamus; MB, midbrain; CB, cerebellum; HB, hindbrain; FT, fiber tract. **b** Spatial mapping of major anatomical regions across all samples. Scale bars, 1 mm. **c** Matrix plot showing mean expression of subregion marker genes combined with all samples. **d** Spatial mapping of cell type annotations across all samples. Scale bars, 1 mm. **e** Cell type composition within each major anatomical region across all samples. Bars represent relative proportions of each cell type.

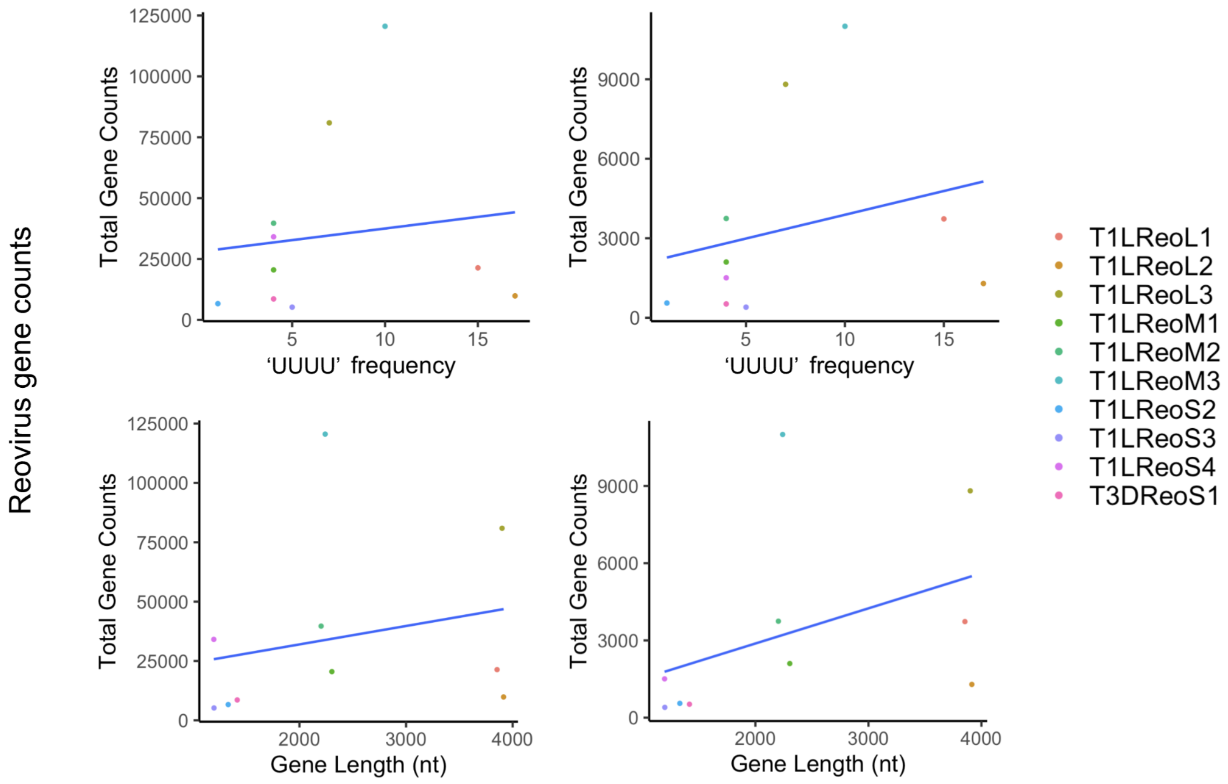


**Supplementary Fig. 2. Expression of the 10 reovirus RNAs does not correlate with gene length or length of four poly-U stretches in the gene.** Across infected 7 DPI replicate 2 (left) and infected 7 DPI replicate 1 (right), the relative number of counts of each of the 10 reovirus RNAs is preserved. However, there is no correlation between gene length and total gene counts for the 10 reovirus genes. Similarly, there is no correlation between the number of UUUU stretches in the gene and total gene counts for the 10 reovirus genes in the 7 DPI (high) sample.

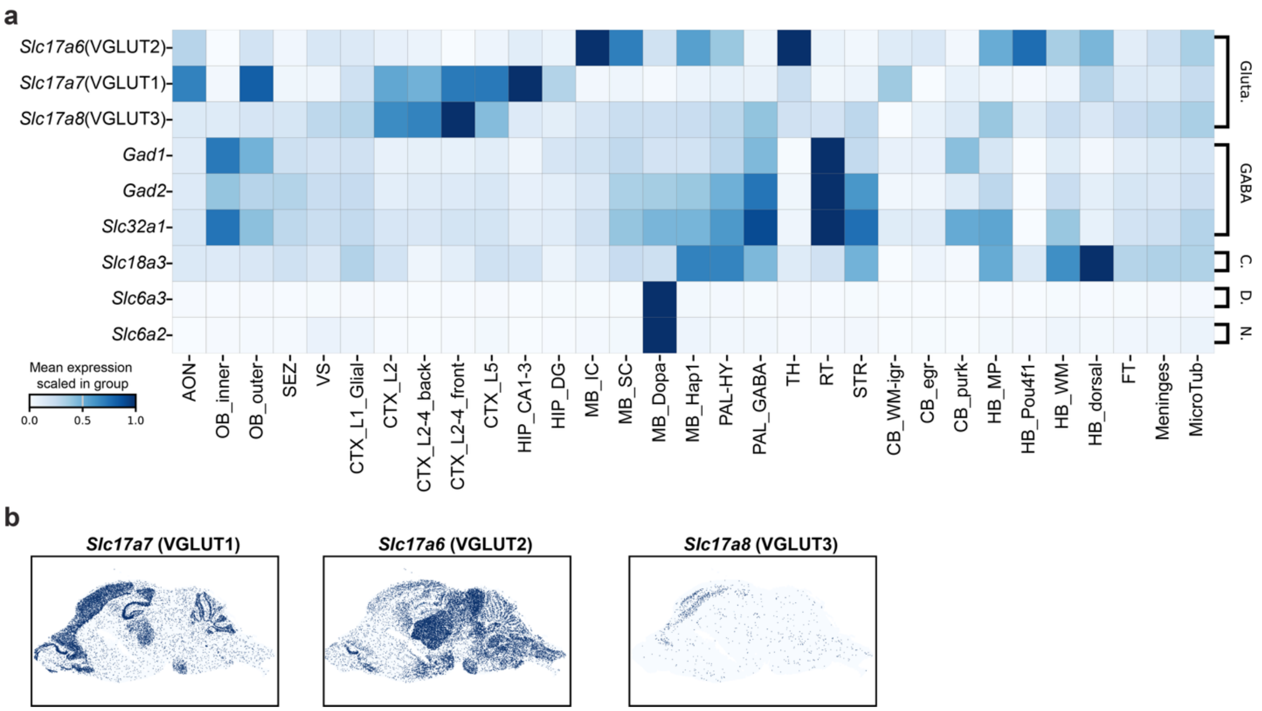


**Supplementary Fig. 3. a** Matrix plot showing mean expression levels for neuronal subtype markers across anatomical subregions. Gluta., glutamatergic; GABA, GABAergic; C., cholinergic; D., dopaminergic; N., noradrenergic. **b** Spatial distribution of VGLUT1-3 in the mock control brain.

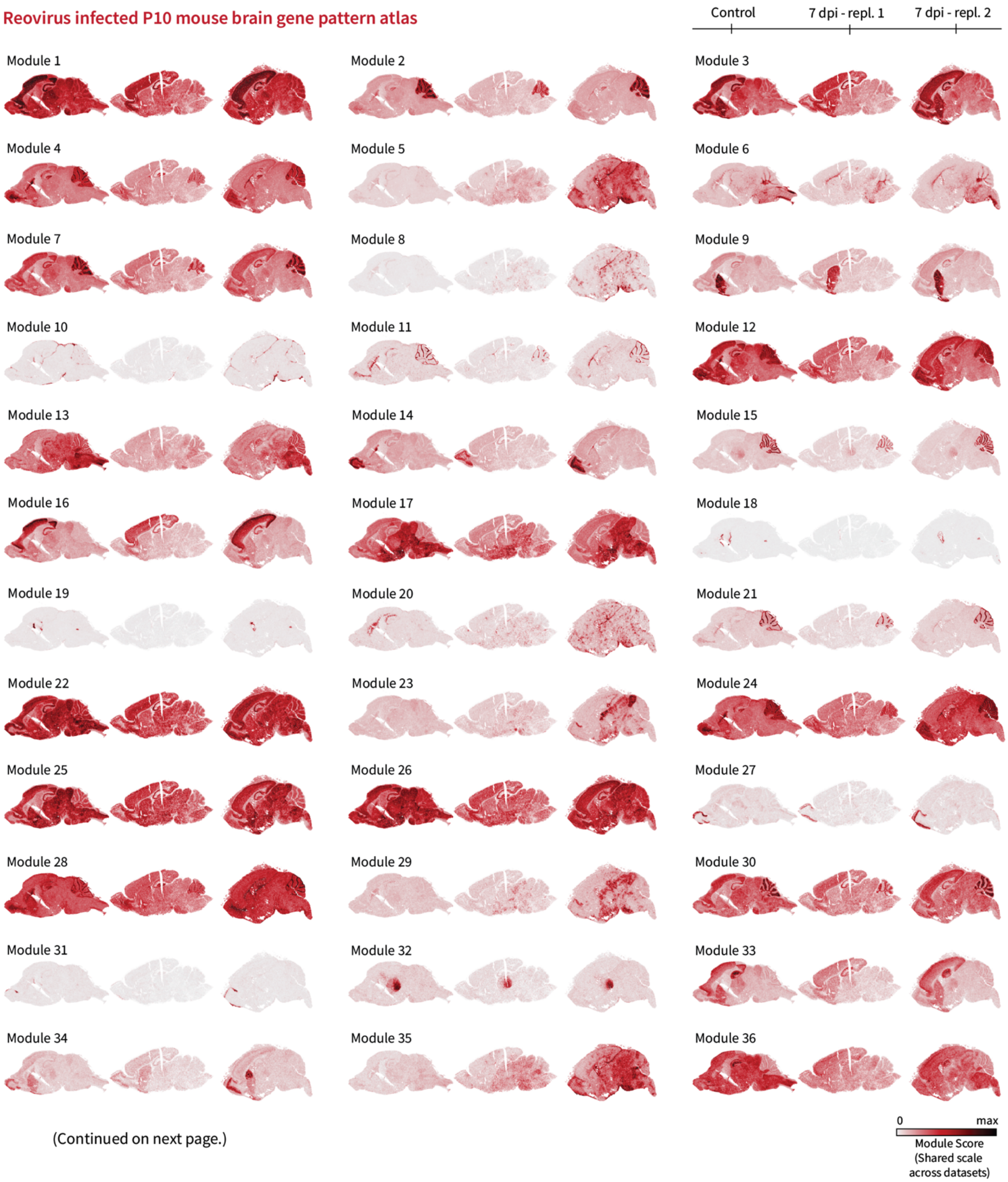


(Continued on next page.)

47
48

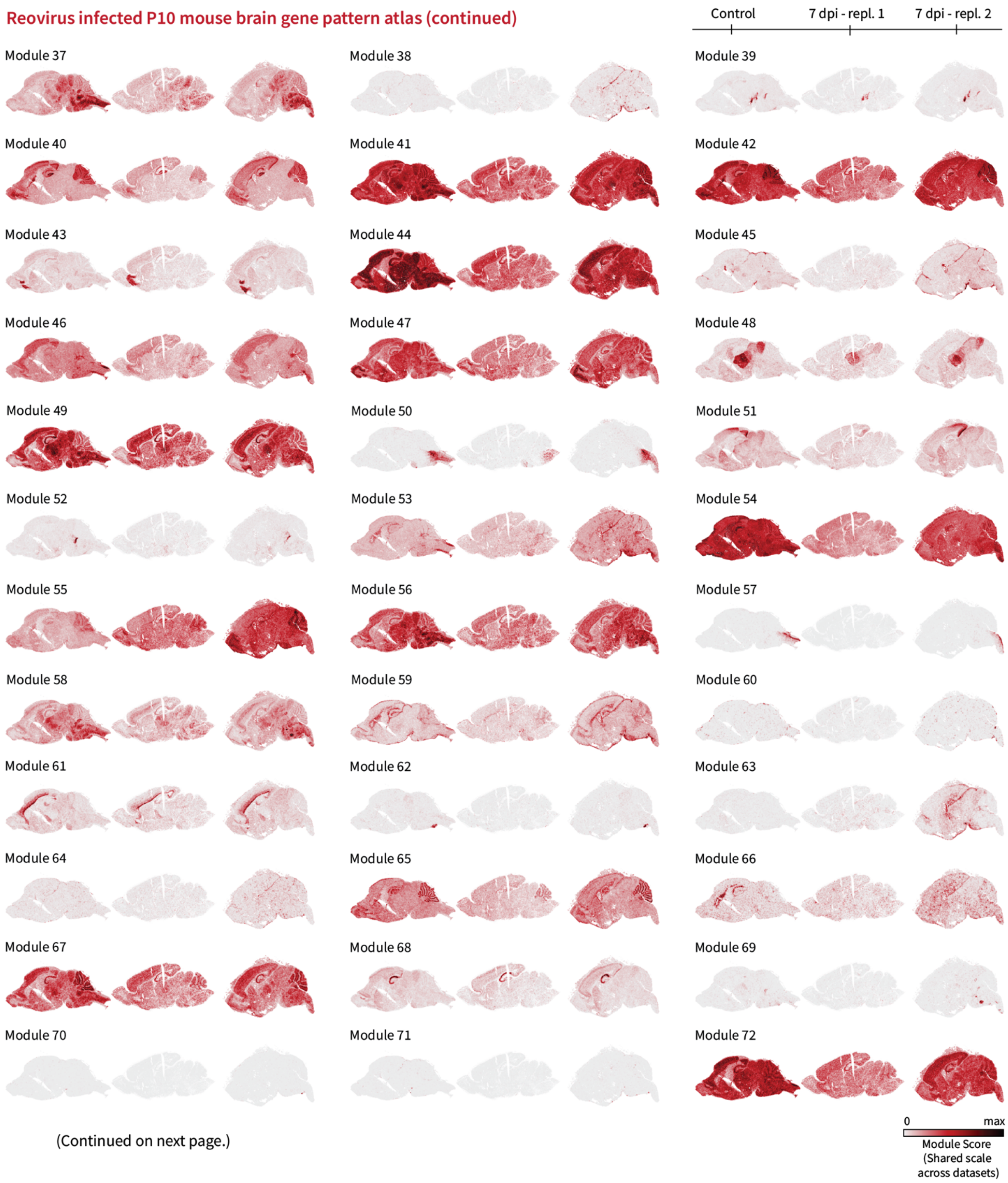


(Continued on next page.)

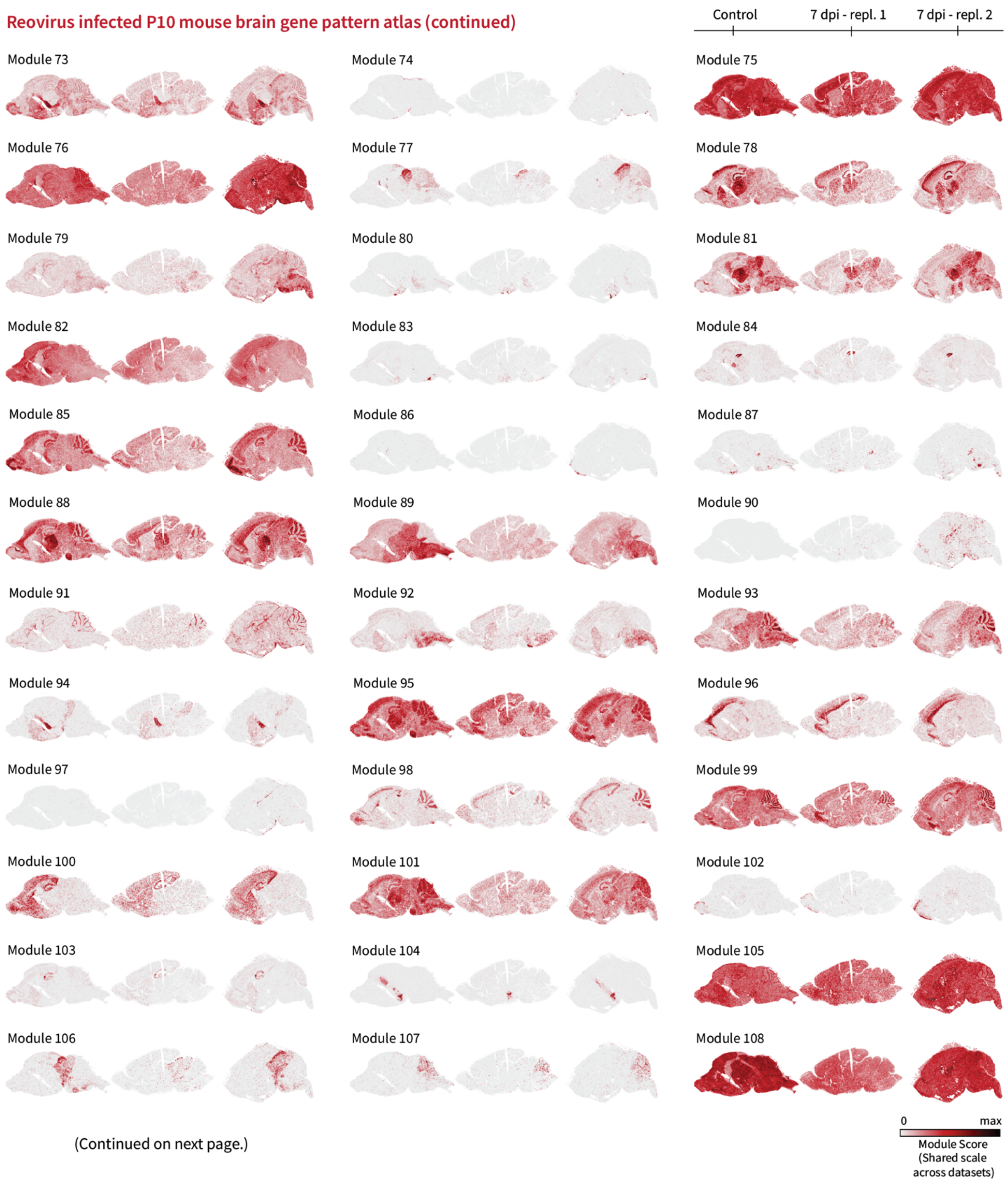
Reovirus infected P10 mouse brain gene pattern atlas (continued)
Control
7 dpi - repl. 1
7 dpi - repl. 2
Module 73
Module 74
Module 75
Module 76
Module 77
Module 78
Module 79
Module 80
Module 81
Module 82
Module 83
Module 84
Module 85
Module 86
Module 87
Module 88
Module 89
Module 90
Module 91
Module 92
Module 93
Module 94
Module 95
Module 96
Module 97
Module 98
Module 99
Module 100
Module 101
Module 102
Module 103
Module 104
Module 105
Module 106
Module 107
Module 108
(Continued on next page.)
0
max
Module Score
(Shared scale
across datasets)

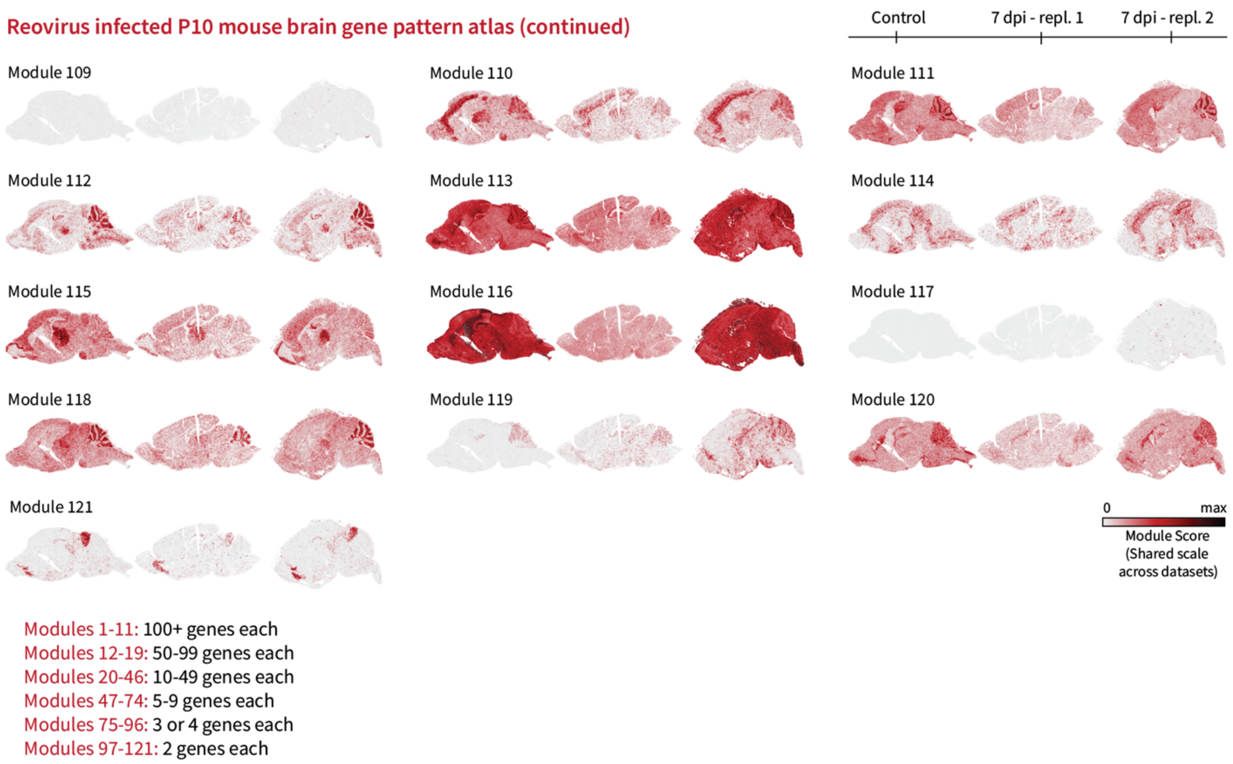


**Supplementary Fig. 4. Gene pattern atlas for 7 day post-infection (dpi) reovirus and control mouse brains.** A total of 121 gene modules were identified using the Smoothie gene co-expression network pipeline. Modules are sorted from largest to smallest. Full gene lists are included in Supplemental Table S1. Module scores are shared across datasets. The score reflects transcript averages across spatial spots after normalizing the transcript of each gene by its maximum value across all datasets.

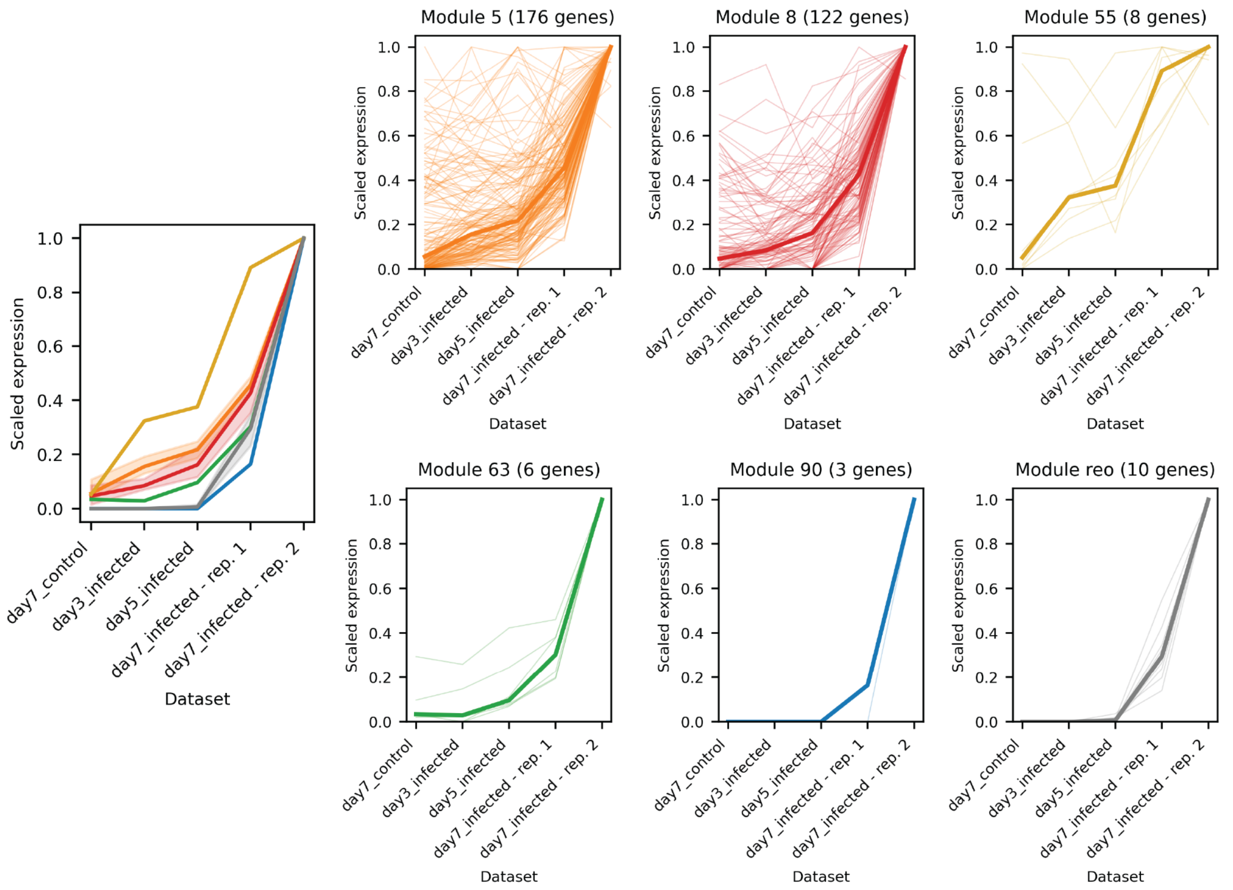


**Supplementary Fig. 5. Relative bulk expression plots for different dynamic gene modules across samples.** Dark lines show the median transcript value across samples. Scaled expression for gene *X* in dataset *D* reflects (gene *X* UMIs / total dataset *D* UMIs), linearly rescaled to make the dataset with the largest value equal to 1.0.

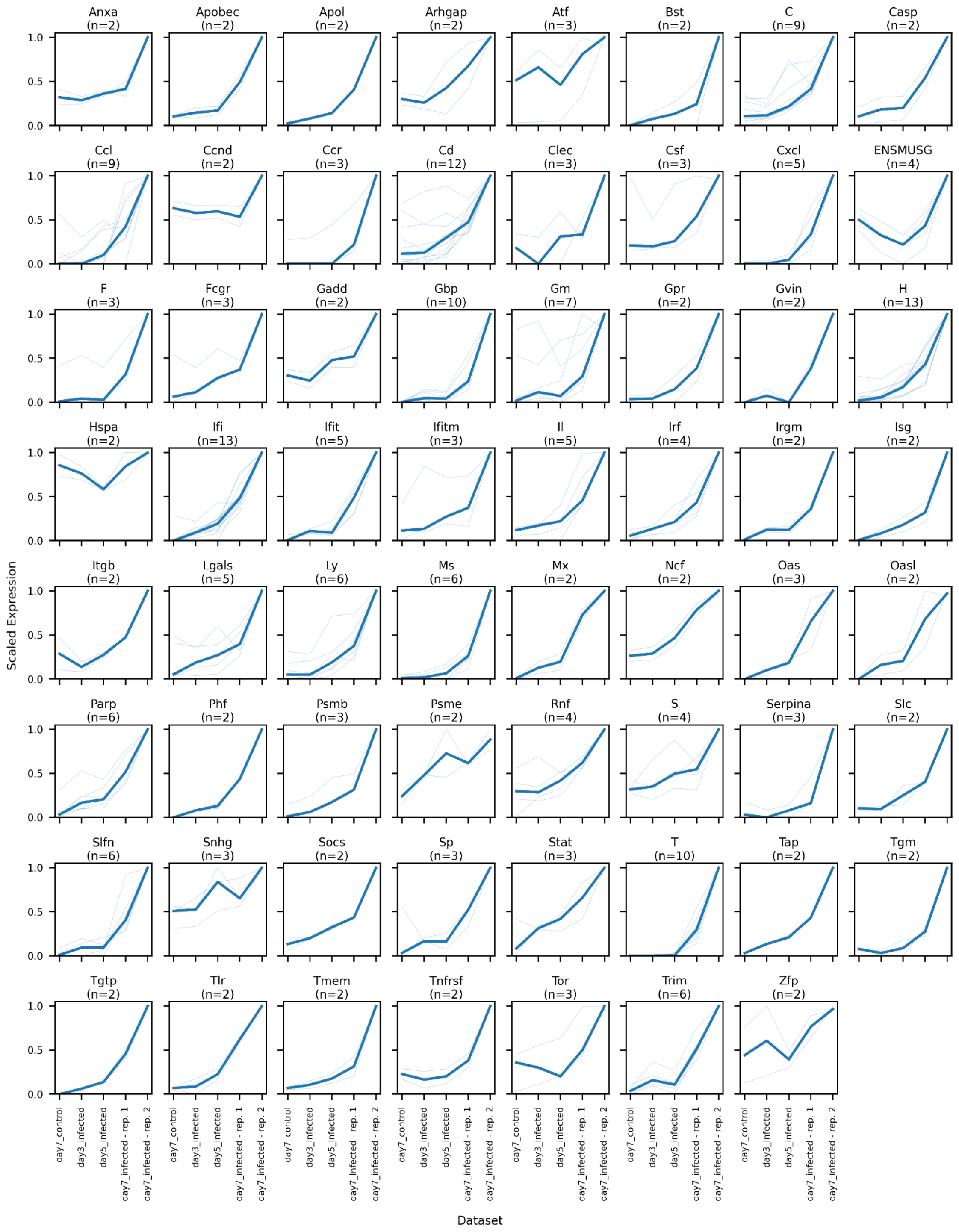


**Supplementary Fig. 6.** Gene family relative bulk expression trajectories across reovirus infection timepoints. Genes are organized by gene name prefix (which often corresponds with meaningful gene family groupings). Only genes that appeared in a dynamic infection response module are included in these plots. Dark blue lines show the median value across samples. Scaled expression for gene *X* in dataset *D* reflects (gene *X* UMIs / total dataset *D* UMIs), linearly rescaled to make the dataset with the largest value equal to 1.0.
